# Experimental investigation using micro-PIV of the effect of microRNA-1 deficiency on motility in *Caenorhabditis elegans*

**DOI:** 10.1101/2020.06.16.119396

**Authors:** S. Ravikumar, M. Fedrizzi, R. Prabhakar, R. Pocock, M. K. O’Bryan, J. Soria

## Abstract

*Caenorhabditis elegans* is a microscopic nematode used extensively as a model organism in studies of neuromuscular function and neurodegenerative disorders. A mutation in *mir-1* affects signalling at the neuromuscular junction. We investigate the effect of this mutation on the propulsive power exerted by nematodes as they grow in size with age. We compare the motility of wild-type and *mir-1(gk276)* mutant nematodes in a Newtonian fluid using a two-component, two dimensional (2C-2D) Digital Microscopic Particle Image Velocimetry (*µ*-PIV) technique. Beating amplitudes of the head and tail, the wavelength of undulatory waves and the swimming speed scale linearly with size in both the wild-type and mutant strains. The beating frequency is independent of size or position along the body. Differences in the magnitudes of these kinematic parameters between the two strains, however, grow systematically with age. The swimming speed scales linearly with the wave speed of the neuromuscular undulation in both nematode strains with a conserved ratio. The magnitude of mean power and mean local fluid circulation in the mutant is significantly lower compared to those of the wild-type animals of the same age. This indicates that a mutation in *mir-1* adversely affects motility in *C. elegans*.

## 1 Introduction

*Caenorhabditis elegans* (*C*.*elegans*) is a multicellular, free-living micro-organism found in rotting organic matter. It is extensively used as a model organism for biomedical investigations and genetic studies related to neuromuscular damage, muscular disorders and ageing. The genome of *C. elegans* has been completely sequenced, and it is estimated that humans and *C*.*elegans* share 60-80% of their genes [Harris et al., 2004]. A variety of mutant strains that over-express or lack a specific gene function have been developed and extensively studied, and comprehensive information on the biological aspects of *C*.*elegans* including their phenotype and genotype is now available [Chen et al., 2005]. In the last few decades, *C*.*elegans* has emerged as a research tool in biomedical studies related to human diseases and it has been estimated that nearly 40% of the genes related to human diseases have an analogue in *C*.*elegans* [Hariharan et al., 2003, Maria and Nektarios, 2010].

Despite these efforts, the crucial question of how growth affects the motility has not been answered. Motility phenotyping of mutant strains with genetic defects similar to age-related diseases in human beings such as Parkinson’s, Alzheimer’s and Huntington’s disease will be useful in biomedical research, particularly for drug development and drug screening applications. In this paper, we quantify and understand the swimming behaviour of *C*.*elegans* during each stage of their growth in a Newtonian fluid medium.

Tavernarakis et al. [1997] investigated the effect of a mutation in *unc-8* on the crawling gait in *C*.*elegans*. Such gain-of-function mutations are known to induce neuronal swelling and severe uncoordination. They showed that the *unc-8* mutation leads to an irregular gait with reduced beating amplitude and wavelength of propagating waveforms from head to tail. On the other hand, a mutation in *trp-4* leads to a defect in a mechanosensitive channel, causing an increased beating amplitude and crawling speed [Li et al., 2006]. Karbowski et al. [2006] investigated mutations affecting *Caenorhabditis* muscles and their effects on parameters in crawling motility such as the velocity, beating frequency and amplitude. They showed that, while some parameters such as crawling velocity, amplitude and wavelength systematically vary with age and are strongly influenced by mutations, other parameters, such as the normalized undulatory wavelength, are conserved across individuals, mutants and even species.

Similar observations have been reported on swimming motility in *C. elegans*. Korta et al. [2007] investigated the effect of mutations in *mec-4* (*d*) and *mec-6* (*u450*) in fluids with viscosity ranging from 0.05-50 Pa s. They showed that, in both the wild-type and mutants, the wavelength of the swimming gait remains unaffected by even a 1000-fold increase in the viscosity, whereas the undulating frequency decreases nonlinearly, as a power-law. The mutants exhibited a higher frequency than wild-type nematodes. Similar observations on swimming kinematics were reported by other studies [Sznitman et al., 2010a, Backholm et al., 2015]. Kinematics have also been studied in

An important advantage in studying swimming is that frictional forces and the mechanical power expended by a microswimmer against those forces can be calculated using hydrodynamic models. The undulatory waveforms in *C*.*elegans* enable it to swim in Newtonian fluids. The low Reynolds numbers corresponding to such motion means that fluid as well as swimmer inertia are unimportant and fluid friction is the dominant force on microswimmers [Purcell, 1977]. Backholm et al. [2015] investigated measured the effect of fluid viscosity on the kinematics of undulation in nematode tethered at their tails to a micro-pipette. They further used Resistive Force Theory [Gray and Hancock, 1955, Lighthill, 1976] to calculate the frictional forces and power exerted by the nematodes in sustaining the undulatory motion. It was observed that the power exerted by the nematodes is constant irrespective of the range of viscosity levels considered in their study. This lead to the conclusion that nematodes modify gait characteristics in response to changes in environmental conditions while keeping the power output constant.

Resistive Force Theory is based on local resistance coefficients that allow the approximate calculation of the frictional force on a segment of the nematode body from its instantaneous velocity obtained by automated image analysis [Sznitman et al., 2010d]. A more direct approach is to measure the fluid velocity field around a swimmer using particle tracking or particle image velocimetry and then calculate the viscous power dissipation in the fluid without depending on approximate resistance coefficients [Sznitman et al., 2010c, Krajacic et al., 2012, Johnson et al., 2016]. Gagnon and Arratia [2016] used the *µ*-PIV technique to analyze flows around nematodes in shear-thinning fluids [Gagnon and Arratia, 2016] and found that the power exerted by an adult nematode in a shear thinning fluid is less than or equal to that exerted in a Newtonian fluids and depended on the effective viscosity of the fluid. Recently, Wan et al. [2014] used the *µ*-PIV technique to compare the swimming behaviour of the the wild-type nematode with that of the mutant strain KG532 with a hyperactive phenotype and a defective gene *kin*-2. They observed that, due to hyperactivity, swimming speed and power exerted are significantly higher in the mutant strain than in wild-type nematodes.

Measurements of the velocity field also reveals the evolution of prominent flow structures around swimming bodies. For instance, Gagnon et al. [2014] showed in xanthum gum solutions in M9 buffer, that shear-thinning fluids does not alter the kinematics of nematode undulation, but the magnitude of the local circulation around the nematode and fluid velocities near the tail portion were found to be significantly enhanced.

Here, we use these powerful techniques to investigate the effect of a mutation in the gene *mir-1* on swimming motility in a Newtonian fluid. The microRNA miR-1 controls neurotransmitter release at the neuromuscular junction [Simon et al., 2008]. *mir-1* also controls the synaptic proteins neuroligin and neurexin [Hu et al., 2012], which in humans, are linked with synaptic function defects associated with autism spectrum disorders [Persico and Napolioni, 2013]. Image analysis is used to characterize the kinematics of nematode motion. We further obtain the mechanical power dissipated and fluid circulation from 2C-2D *µ*-PIV measurements of the two in-plane components of velocity fields around swimming nematodes as a function of position in the focal plane. We track changes in these motility characteristics with age as nematodes progress through their first three larval stages of growth and increase their length from 220 to 490 *µm*.

## 2 Measurement system

### 2.1 Nematode preparation

Wild-type (N2) and mutant (mir-1(gk 276)) C. elegans were used for this experimental investigation. N2 worms are used as the reference behaviour in different stages of their growth.

Nematodes were grown on agar plates (Nematode Growth Medium) [Brenner, 1974] in an incubator maintained at a temperature of 20*°*C and well fed with *E. coli* (OP50 strain). Animals were synchronized by washing a mixed-stage sample with M9 buffer and the eggs were collected and transferred to NGM plates (at least six plates were prepared for each strain) seeded with *E. coli*. The plates were kept in the incubator and washed at different time intervals to sample nematodes of different sizes. A total of 65 to 70 individual nematode samples were investigated for each kind. Throughout the experiment, nematodes were maintained at a room temperature between 17*°*C to 20*°*C, and the total duration of one complete experiment of 4 to 5 nematode samples was less than 5 to 10 minutes.

### 2.2 2C-2D *µ*-PIV system

An in-house 2C-2D *µ*-PIV system was used to investigate the swimming behaviour of the wildtype and mutant nematodes. The 2C-2D *µ*-PIV experimental set-up consists of: (*i*) a pulsed LED light source to illuminate the flow field, (*ii*) a microscope objective with 40x magnification and the optical sub-systems (lens, mirror, etc.) to focus the nematode along with the flow tracer particles, (*iii*) a digital 5 MPx sCMOS camera to acquire the single-exposed images, (*iv*) a Beagle Bone Black programmable real-time controller [Fedrizzi and Soria, 2015] used to synchronised the sCMOS exposure with the pulsed LED light source and (*v*) a fluid chamber. Figure 1 shows the schematic representation of the experimental set-up and the apparatus used, which was fixed on an optical table. The camera and the microscope objective were positioned vertically above the fluid chamber as shown in figure 1. The fluid chamber was fixed horizontally on top of an optical x-y traverse that allowed optical access and accurate positioning of the fluid chamber. The LED and lens were positioned horizontally on the optical table on optical rails and aligned in-line with the microscope objective and the camera. A 45-degree mirror was used to direct the LED light vertically up into the fluid chamber through its glass bottom surface. The mirror was positioned at the bottom of the mount and adjusted such that the required illumination intensity was achieved. The light passes through the lens and mirror which expands and deflects the light beam used to illuminate the entire flow.

**Figure 1:**
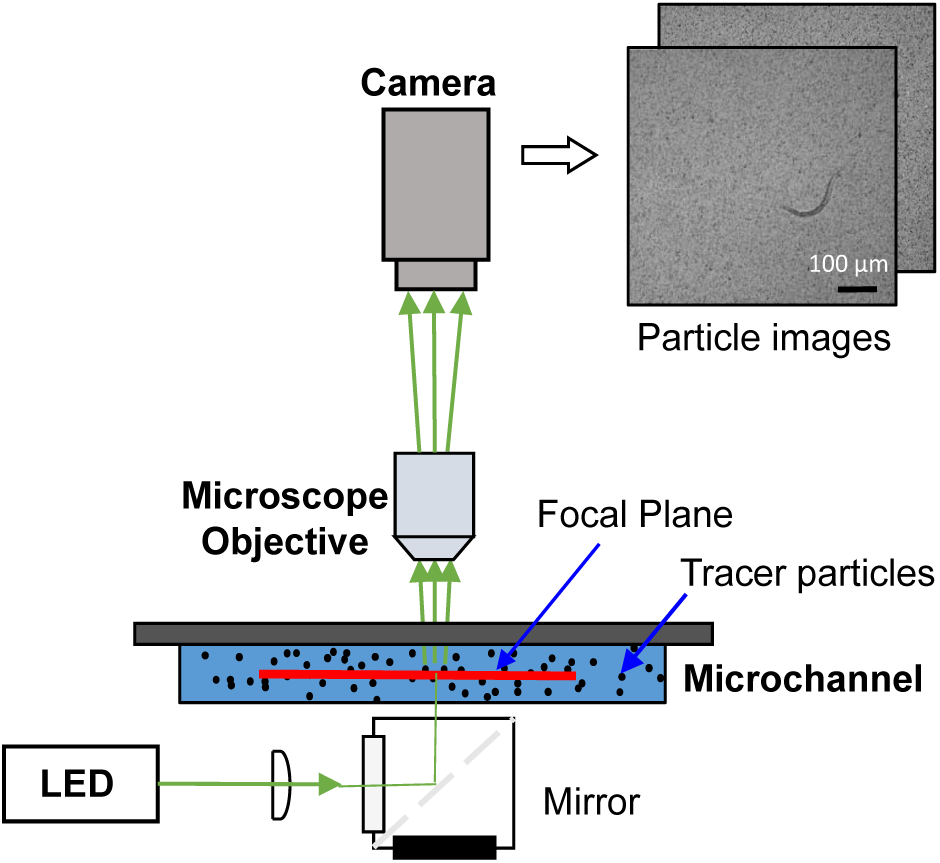
Schematic representation of experimental set-up.

The experiment was performed in Newtonian, water like M9 solution (*ρ* = 10^3^ *kg/m*^3^ and *µ* = 1 *mPa*.*s*). Polystyrene spheres of 1 *µm* in diameter were used as the flow tracers. The concentration and inertia of these particles was not found do alter the swimming behaviour of nematodes in the fluid. Nematodes were diluted with the M9 solution along with the flow tracer particles in the ratio of 1:10, i.e. 0.1% of particles for 1*cc* of the M9 solution to achieve a uniform seeding density. During the experiment, a 0.5 *µl* droplet of the M9 solution along with the tracer particles and nematodes was transferred to the fluid chamber. The fluid chamber was made up of two glass microscope slides sandwiched together and separated by a distance (*H*) using the adhesive tape. The gap between the glass slides was maintained at 50-60 *µm* throughout the experiment since the stage 3 nematodes can grow up to an average diameter of 30 *µm* and it was found that this depth is sufficient for nematodes in the first three stages to swim freely in the observation plane which was in focus. Since the experiment involves living micro-organism, to avoid the sedimentation of particles and the accumulation of biological waste which may alter the swimming behaviour, the nematodes were mixed with the M9 solution and particles just before the start of each experiment. Thus, the particle mixture was prepared for each sampling and flushed after each experiment.

A sequence of 75 image pairs with a temporal resolution of Δt = 15 ms or 20 ms between each pair was acquired at a rate of 15 pairs per second and at least 4 to 5 swimming cycles were recorded to maximise the resolution and accuracy of kinematic measurements. The time interval between the image pairs was selected to ensure that the maximum tracer particle displacement was 6-8 px. A total of 65 to 70 independent nematode samples of each kind were imaged using this methodology. During the image acquisition, the middle of the fluid chamber was set to be the focal plane to avoid wall effects. Out of plane recordings and the images with more than one nematode were discarded. Table 1 shows a summary of recording parameters and the resolution of the 2C-2D *µ*-PIV system used in this study.

**Table 1:** Summary of relevant parameters of *µ*-PIV measurement.

|  | Particulars | Value/Type | Unit |
| --- | --- | --- | --- |
| **Seeding** | Particle Type | Spherical |  |
| | Diameter | 1 | $\mu m$ |
| **Imaging** | Camera Type | sCMOS |  |
| | Resolution | 2560x2152 | $px$ |
| | Image pair acquisition frequency | 30 | $Hz$ |
| | Total acquisition duration | 2.6 | $s$ |
|  | Total image pairs per sample | 80 |  |
| | Time interval between images | 15-20 | $ms$ |
| | Maximum particle displacement | 6-8 | $px$ |
| | Pixel size in object space | 0.3 | $\mu m/px$ |
| **Optical system** | Numerical aperture | 0.65 |  |
|  | Magnification | 40x |  |
| | Depth of field/focus | 1.4 | $\mu m$ |

### 2.3 Kinematic measurements

The primary kinematic data of the wild-type and mutant nematodes for the kinematic study were obtained by tracking the nematode body positions and the corresponding coordinates in each instantaneous time frame which is approximated by the centreline (skeleton) of the body. To segment the nematode body portion without the fluid particle tracers from the raw image and extract the nematode body centreline, a number of morphological image processing operations were performed including dilation, fill holes, erosion and skeletonisation [van den Boomgard and van Balen, 1992, Ongwattanakul et al., 2003, Soille, 1999]. Initially, the dilation, erosion and fill holes operations were performed to remove the particle traces from the background and segment the nematode body shape accurately. Finally, the skeletonisation [Lam et al., 1992, Soille, 1999] was performed on the binary images of the nematode. This process involves removal of pixels along the surface of the nematode’s body without breaking the nematode’s body apart such that the pixels in the centre of the body represents the skeleton. After extracting of the skeleton, a smooth polynomial curve fit was established to approximate the centre line of the nematode body posture at any given instant of time. These curves were parametrised to obtain uniformly distributed points (N = 50) along the centreline and represent a continuous 2D curve which is differentiable. Then length of the nematode was obtained by measuring the length of the body centreline. The length of the body centreline is defined as the sum of the distances between the body coordinates (*s*) distributed along the nematodes body with each point to point distance calculated as 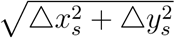 [Cronin et al., 2005], where *x*_*s*_, *y*_*s*_ are the corresponding coordinates. The kinematic parameters such as the bending curvature values (*k*) along the nematodes body, beating frequency, amplitude and wavelength were measured using these body coordinates.

#### Spatio-temporal evolution of swimming gait

*C.elegans* are undulatory swimmers with sinusoidal bending waveforms which propagate from head to tail [Korta et al., 2007, Sznitman et al., 2010b,c]. Hence, the swimming behaviour was characterised by analysing the bending curvature of the swimming gait (which evolves in space and time) in each stage of their growth. The coordinates along the body, *s* is used to define the bending curvature along the nematode’s body which is given by the equation,

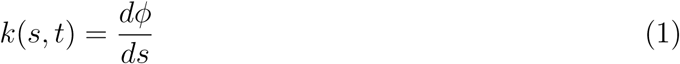

where *ϕ* is the angle made by the tangent drawn at each point along the centreline of the body to a fixed frame of reference (x-axis) and *s* is the coordinates along the body spanning from the head (*s = 0*) to tail (*s = L*) [Korta et al., 2007, Sznitman et al., 2010b,a,c, Krajacic et al., 2012, Shen et al., 2012, Gagnon et al., 2014].

#### Frequency of beating

The beating frequency (*f*) of both wild-type and mutant nematodes was obtained by extracting the curvature values *k* (*s, t*) from the well-defined curvature fields (spatiotemporal evolution of the bending curvature) at distinct body positions spanning from head to tail (*s/L* values of 0.2, 0.5 and 0.8). The beating frequency is defined as

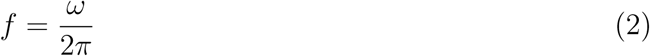

where *ω* is the angular frequency (*rad/s*) computed by extracting the curvature values at distinct body positions along the nematode’s body and computing the one dimensional Fourier transform [Korta et al., 2007, Sznitman et al., 2010a,b, Krajacic et al., 2012, Shen et al., 2012].

#### Amplitude of beating

The beating amplitude (*A*) at different body positions was obtained by tracing the trajectories delineated by the body centreline coordinates (*s*) at distinct body positions (*s/L*=0.2, 0.5, 0.8) over multiple beating cycles (≈4-5) and then computing the one dimensional fast Fourier transform (FFT). The coordinate positions traced in space over swimming multiple cycles as a function of time is used as inputs to the FFT operation. Then the head beating amplitude is obtained by taking the absolute value of complex amplitude output from the FFT operation, which is given by *A*_*h*_ = 2*(abs(*FFT* (*H*_*c*_)/*N*)), where *H*_*c*_ is the head coordinate traced over multiple beating cycle in each time step and *N* is the number of sampling points. Since the output is a double sided spectrum, the output amplitude is multiplied by a factor of 2. Similarly, the body beating amplitude (*A*_*b*_) and tail beating amplitude (*A*_*t*_) is obtained by tracing the corresponding coordinates in space and time.

#### Swimming speed and wavelength

The centroid coordinates (*c*_*c*_) of the nematodes were obtained at each instant of time by tracing the coordinates (*x*_*c*_, *y*_*c*_) of the local centroid of nematode body postures in each time frame. The swimming speed of both kinds of nematodes was obtained by differentiating these centroid positions with respect to time which is given by the equation

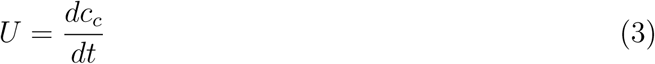

where *c*_*c*_ is the centroid coordinates of the nematodes body traced over multiple swimming cycles. Another kinematic parameter is the wavelength (*λ*) which is one of the main characteristics of the propagating waves travelling along their body from head to tail. The wavelength is obtained by computing the wave speed (*c*) in each stage of the nematodes growth. The slope of the diagonal lines in the curvature fields (*k* (*s,t*)) gives the wave speed (as seen in figure 6 (*e*)). Then the wavelength is computed using the equation,

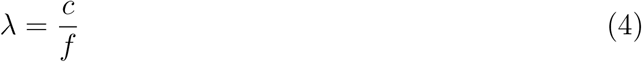

where *f* is the beating frequency (rate of occurrence of the propagating waveforms).

**Figure 2:**
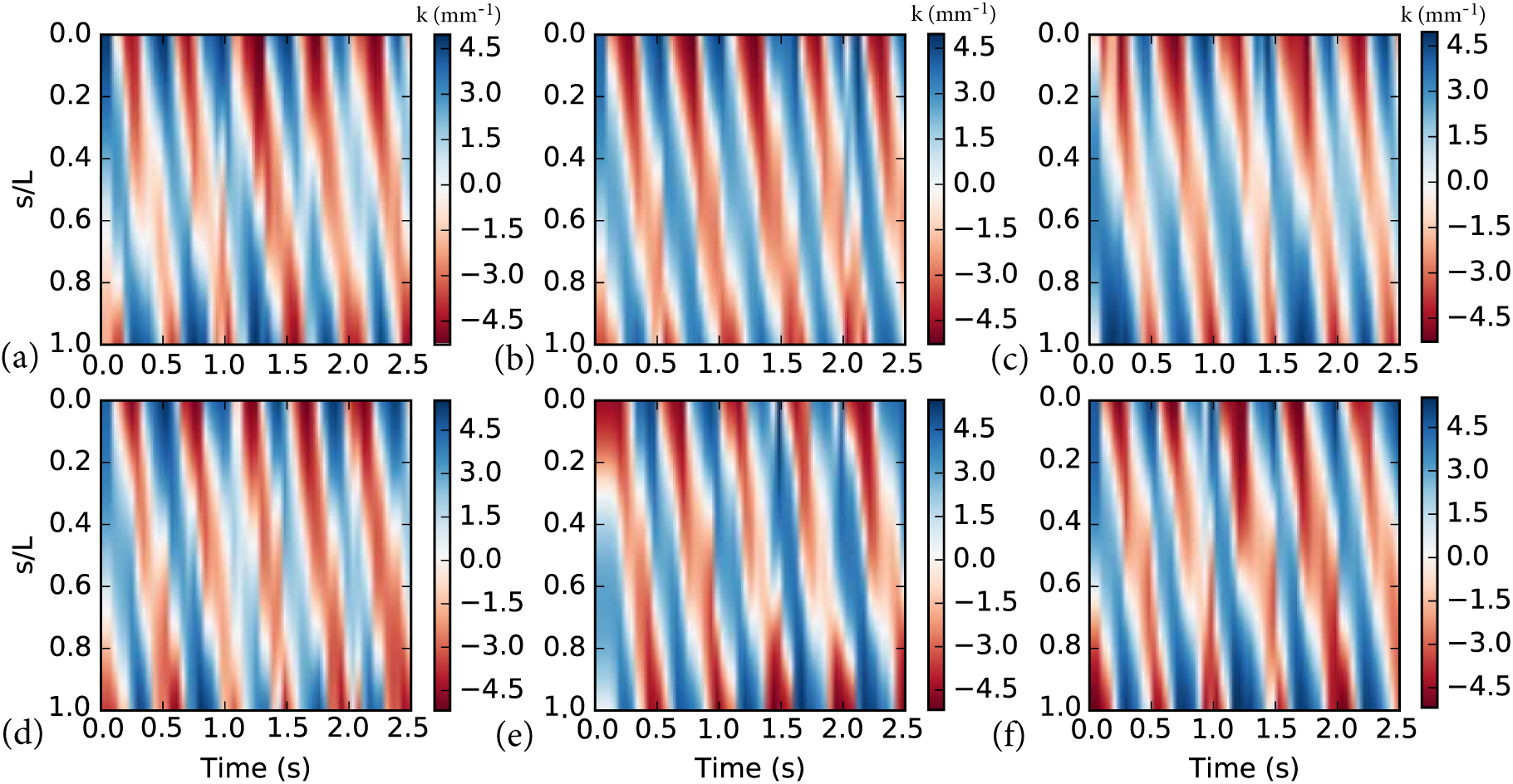
Spatiotemporal evolution of the bending curvature (*k*) of (*a, b, c*) wild-type and (*d, e, f*) mutant nematode samples in each stage (1, 2, 3) of their growth. Alternate colour code refers to the positive and negative values of ‘*k*’ respectively.

**Figure 3:**
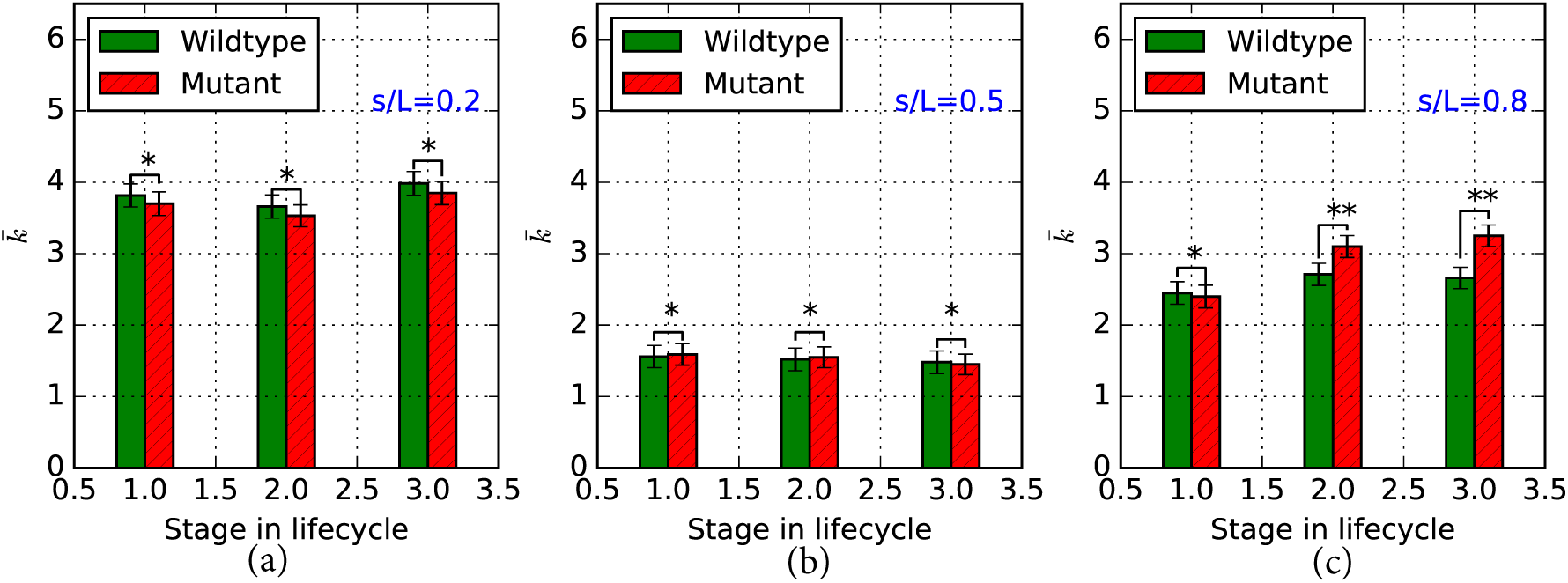
(*a,b,c*) Time averaged 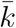 of the wild-type and mutant nematodes at different body positions (*s/L* = 0.2, 0.5, 0.8) in each stage of their growth. Error bar represents the standard deviation. * denotes no significant difference and ** denotes significant difference based on t-test for p <0.05.

**Figure 4:**
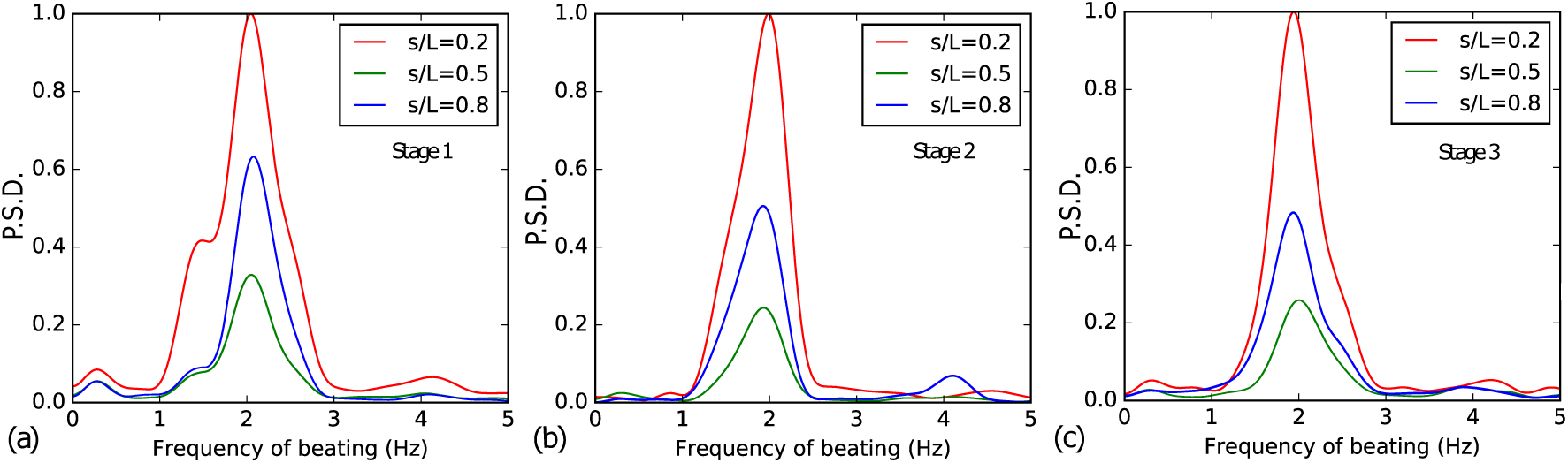
(*a,b,c*) Normalized mean frequency spectrum of wild-type nematodes at different body positions (*s/L* = 0.2, 0.5, 0.8) in each stage of their growth.

**Figure 5:**
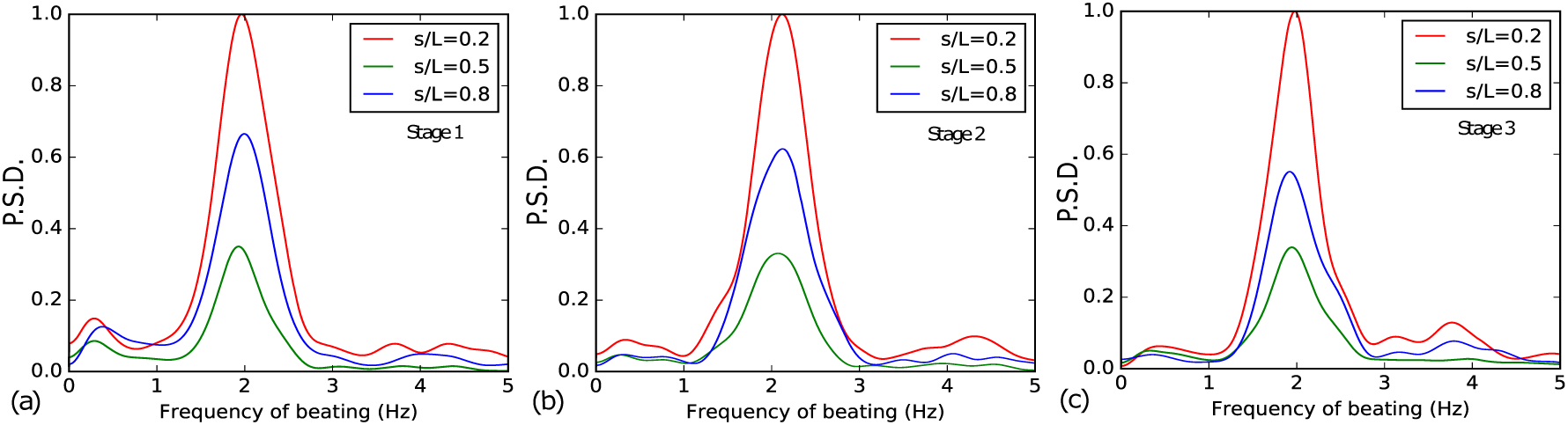
(*a,b,c*) Normalized mean frequency spectrum of mutant nematodes at different body positions (*s/L* = 0.2, 0.5, 0.8) in each stage of their growth.

**Figure 6:**
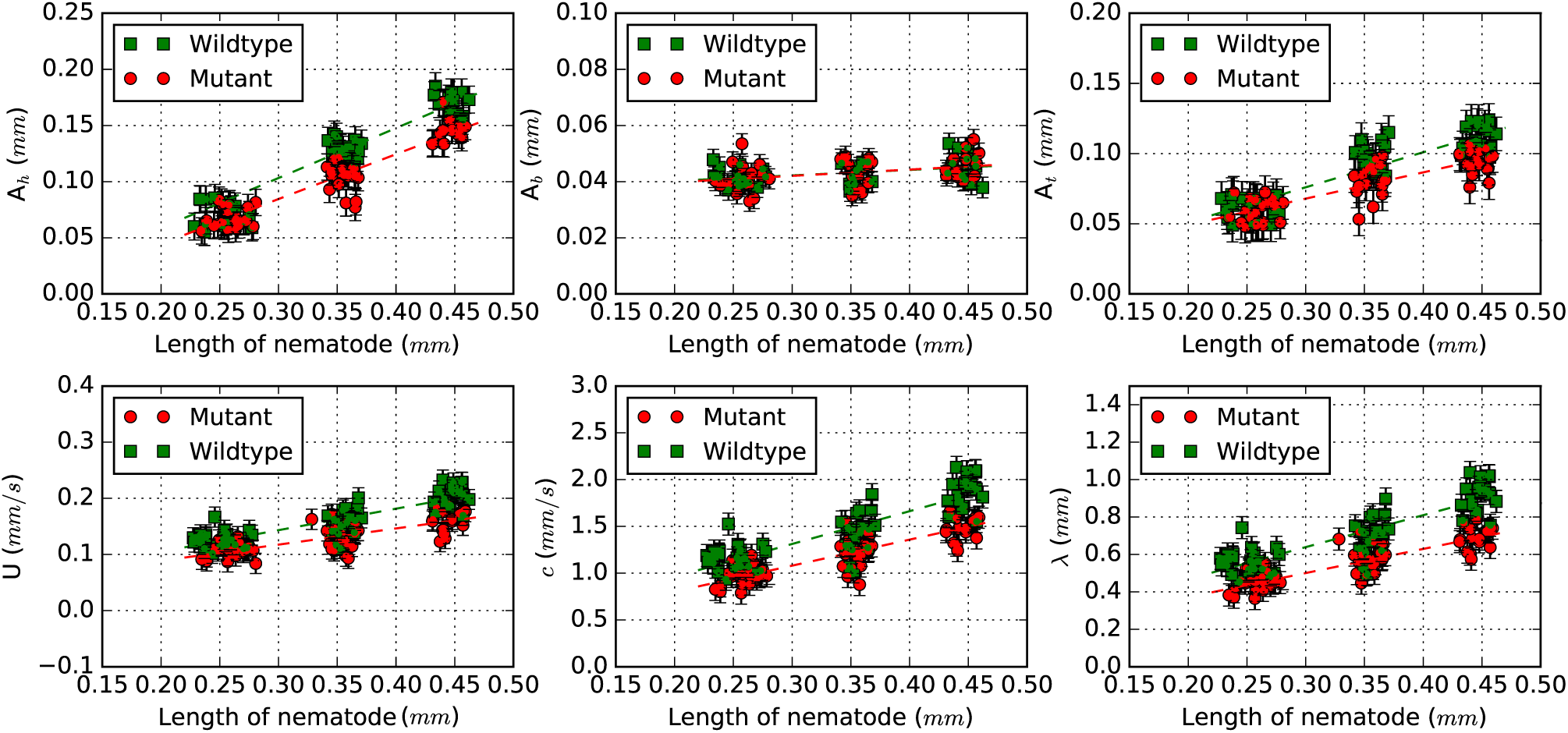
(*a*) Head beating amplitude (*A*_*h*_) at *s/L*=0.2. (*b*) Body beating amplitude (*A*_*b*_)at *s/L*=0.5. (*c*) Tail beating amplitude (*A*_*t*_) at *s/L*=0.8. (*d*) Swimming speed (*U*). (*e*) Wave speed (*c*). (*f*) Wave length (*λ*). The dashed lines represent the linear scaling of the measured values with the size (*L*). Error bar represents the standard error across samples. The least square fit to the kinematic data yields *A*_*h*_ = 0.49*L*-0.043 (wild-type) and *A*_*h*_ = 0.41*L*-0.035 (mutant); *A*_*b*_ = 0.0195*L*+0.036 (wild-type) and *A*_*b*_ = 0.0205*L*+0.035 (mutant); *A*_*t*_ = 0.24*L*+0.010 (wild-type) and *A*_*t*_ = 0.19*L*-0.013 (mutant); *λ* = 1.66*L*+0.11 (wild-type) and *λ* = 1.32*L*-0.12 (mutant).

### 2.4 Fluid velocity measurements

The local instantaneous velocity fields were obtained from the single-exposed image pairs of the fluid tracer particles using the multi-grid/multi-pass cross-correlation digital PIV (MCCDPIV) algorithm introduced by Soria [1996]. Details of the performance, accuracy and uncertainty of the MCCDPIV algorithm with applications to the analysis of single exposed PIV and holographic PIV (HPIV) images have been reported in Soria [1998] and von Ellenrieder et al. [2001], respectively. The MCCDPIV algorithm incorporates the local cross-correlation function multiplication method introduced by Hart [1998] to improve the search for the location of the maximum value of the cross-correlation function. For the sub-pixel peak calculation, a two dimensional Gaussian function model was used to find, in a least square sense, the location of the maximum of the cross-correlation function [Soria, 1994].

Prior to MCCDPIV analysis the background subtracted flow fields without the nematode were segmented into small interrogation regions. Each interrogation region in each image pair was then analyses using MCCDPIV to yield the local average displacement of particles within each interrogation region. The corresponding instantaneous velocity fields were obtained by dividing the local average displacements found in each interrogation region using the known time interval (Δt) between the image pair. Table 2 shows the summary of parameters used for cross-correlation PIV analysis. Two-pass processing was performed on each set of the acquired image pairs. The interrogation window size used in the first pass was 64×64 pixels with a grid spacing of 32×32 pixels, and the window size of 32×32 with a grid size of 16×16 was used in the second pass. The interrogation window size was chosen in such a way that at least 6-8 particles per interrogation window were recorded. The generalised validation was performed using the normalised median filter [Westerweel and Scarano, 2005] and the displacement vectors with large deviations compared to the local average displacement of neighbouring vectors (using 3×3 neighbouring vectors) are identified and eliminated using a suitable cut-off value selected by measuring the residual. The percentage of spurious vectors in any instantaneous velocity field measured was less than 5%.

**Table 2:**
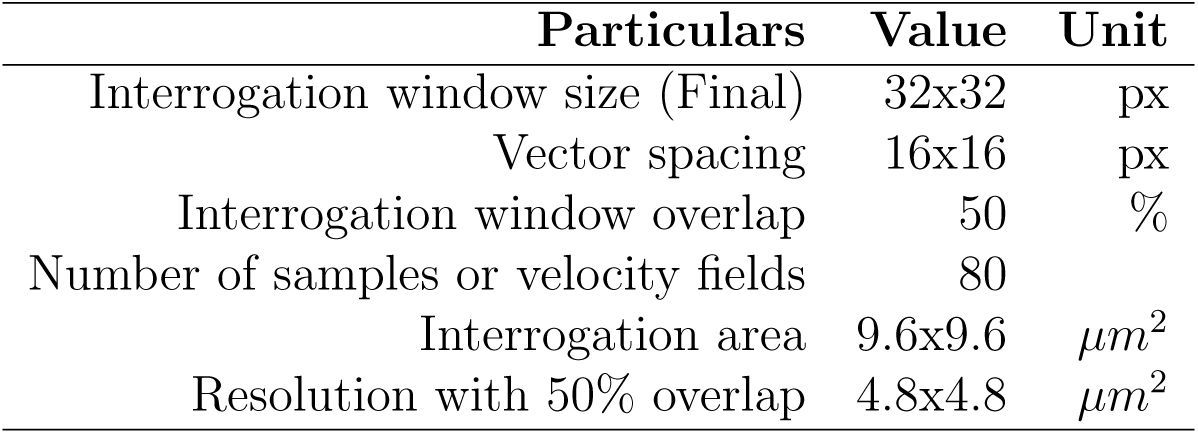
Summary of parameters used for cross-correlation PIV analysis.

#### Power exerted by the nematode

The Reynolds number (*Re*) of the flow generated by the *C.elegans* approximately ranges from 0.04 to 0.1 and almost tends to zero either because of the very small length scales (250 to 460 *µ*m) or very small velocity magnitude generated by these nematodes in the fluid medium. In an incompressible fluid medium, at very low Reynolds number, the effect of inertia and the body forces are negligible with the only dominant force being the viscous stresses and the Stokes equation governing the fluid motion [Purcell, 1977]. In order to satisfy conservation of energy, the energy dissipated by the nematode must be equal to the energy dissipated in the fluid [Purcell, 1977, Gagnon and Arratia, 2016]. In other words, in a control volume, the power exerted by the nematode is equal to the viscous dissipation in the fluid. Thus, the surface integral of the rate of work at the nematodes body surface is converted into a volume integral of viscous dissipation by using Gauss’s divergence theorem assuming that the velocity (*u*) vanishes far away from the nematode’s body surface [Purcell, 1977, Stone and Samuel, 1996, Gagnon and Arratia, 2016]. The power (*P*) exerted in the fluid can be estimated by calculating either the nematodes mechanical power at the surface or the viscous dissipation rate of the surrounding fluid using the equations,

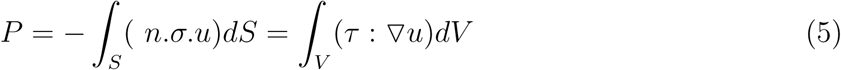

where *n* is the unit normal vector, *u* is the velocity, *σ* is the total stress tensor, *τ* is the shear stress tensor, and the right-hand side of the equation is the viscous dissipation in the fluid (*ϕ*_*v*_). The nematode beats primarily in the observation plane and the particle image velocimetry images for the 2C-2D PIV were recorded in the mid-plane of the nematode. Therefore, in this study, the power exerted by the nematodes was obtained by estimating the viscous dissipation in the fluid using the planar 2C-2D PIV data measured at the centre of the fluid chamber. In this case, the viscous kinetic energy dissipation is given by the equation,

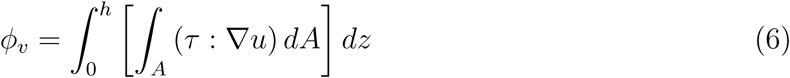

where h is the depth of the flow chamber normal to the 2D velocity measurement plane in the z-direction. At the mid plane (focal plane), where z = 0; the viscous kinetic energy dissipation per unit depth is given by the equations,

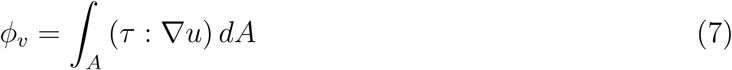

The components of the dissipation function (*ϕ*_*v*_) per unit depth of a Newtonian fluid is given by,

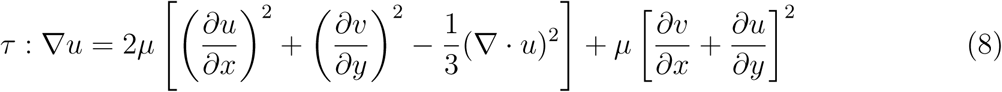

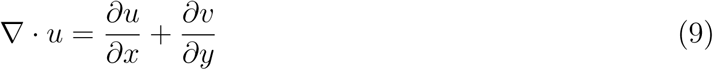

where *µ* is the dynamic viscosity of the fluid and (*u, v*) are in the in-plane fluid velocity components in the (*x, y*) directions respectively. A simple scaling analysis Ravikumar [2018] shows that the bias error in this approximation due to the limited 2C-2D velocity measurements is of the order of 30%. This error can only be avoided by having 3C-3D PIV data, which is exceedingly difficult to obtain with the available experimental infrastructure and is a limitation of the present study. Nevertheless, since the estimated dissipation function (*ϕ*_*v*_) per unit depth is used in a comparative sense only to investigate trends if mutation in mir-1 affects signalling at the neuromuscular junction of *C. elegans*, it is deemed to be a reasonable approach.

## 3 Results and discussion

### 3.1 Spatio-temporal evolution of swimming gait of wild-type and mutant nematodes

Figure 2 (*a,b,c*) shows the spatio-temporal evolution of *k* (*s,t*) of three wild-type nematode samples from the first three stages of their growth. The vertical axis corresponds to the body positions *s/L* (coordinate along the body represented as a fractional distance along the body length). The curvature values are colour coded, and the alternate colours represent the positive and negative values of *k* respectively. The diagonal lines represent the characteristic bending waveforms which propagate from head to tail and correspond to the alternating dorsal and ventral contractions of the nematode body during swimming. These lines travel along the body, and the magnitude of *k* decays from the head (*s* = 0) to tail (*s* = *L*). The slope of these lines represents the wave speed, and their rate of occurrence represents the frequency of beating (*f*) [Korta et al., 2007, Sznitman et al., 2010b,a,c, Krajacic et al., 2012, Gagnon et al., 2014].

It is clear from the figure 2 (*a,b,c*) that the spatial patterns are similar, and the magnitude of *k* (*s,t*) decays from head to tail in each stage of their growth. This observed feature and the curvature values obtained in each stage of their growth are consistent with the results from previous studies (*k*≈ 4-6 *mm*^*−*1^) of the adult wild-type nematodes (≈ 1*mm* in length) [Korta et al., 2007, Sznitman et al., 2010a,c, Gagnon et al., 2014]. No significant difference in the periodic swimming behaviour and the spatio-temporal form of the bending curvature is observed in each stage of their growth, indicating that the swimming gait is highly robust and independent of the nematode size. Figure 2 (*d,e,f*) shows the spatio-temporal evolution of *k* (*s,t*) of three mutant nematode samples from the first three stages of their growth. No significant difference in the rate of occurrence and the periodic spatial patterns of the swimming gait are observed between wild-type and mutant nematodes in each stage of their growth. However, larger values of *k* (*s,t*) are observed at the tail portion of the mutant nematodes in the later stages of their growth. This is clearly visible in figure 2 (*e* and *f*) (dark red colour at the tail portion (*s/L* = 0.8)) where the values of *k* decays from head to mid portion of the body and again increases at the tail portion, whereas a normal decay in *k* is observed among the wild-type nematodes (as shown in figure 2 (*b* and *c*)) of the same size. Also, the variations observed among the mutant nematodes as they mature contrast the characteristic swimming behaviour of the wild-type *C.elegans* of the same size.

Figure 3 (*a,b,c*) show the quantitative comparison of the time averaged curvature values 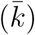 at distinct body positions (*s/L* = 0.2, 0.5, 0.8) of the wild-type and mutant nematodes during their growth. The time averaged mean curvature 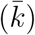 is obtained using the equation,

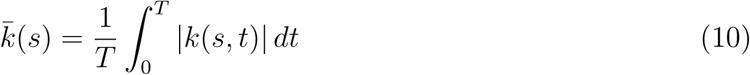

where *T* is the total time period and |*k*(*s,t*) | is the absolute curvature values at different body positions (*s/L*). The average 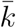 values at the head portion (*s/L* = 0.2) and the mid-body portion (*s/L* = 0.5) of both wild-type and mutant nematodes show a negligible difference in each stage of their growth. However, the average 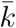 values at the tail portion (*s/L* = 0.8) of the matured mutant nematodes (in stage 2 and stage 3) are higher than that of the wild-type nematodes as they mature, suggesting that the mutant nematode investigated in this study exhibit defect in motion as they mature when compared to the standard assay of the wild-type nematodes of the same size.

### 3.2 Primary kinematic parameters

Figure 4 shows the mean normalised frequency spectrum of *k* (*s,t*) of the wild-type nematodes at different body positions in each stage of their growth. Other discernible peaks as seen in the graph correspond to harmonics in addition to noise in the measurement. A single frequency peak is observed in the frequency spectrum in each stage (S1 ≈ 2.1 ± 0.26 Hz, S2 ≈ 2.03 ± 0.27 Hz, S3 ≈ 1.82 ± 0.29 Hz). Sznitman et al. [2010b] showed that adult wild-type nematodes (≈ 1mm in length) beat at a nearly constant frequency of ≈2 Hz in Newtonian fluids which is similar to the values obtained in this study. The beating frequency is found to be independent of the nematode size and distinct positions along their body, and it is nearly constant (with no significant difference) in each stage. This observation is consistent with the results of Karbowski et al. [2006], where they showed that the temporal form of the crawling gait (on wet surface) was found to be independent of several physical parameters including the size of the wild-type nematodes. Figure 4 also illustrates the normalised curvature amplitudes at different body positions. The maximum amplitude is at the nematodes head portion (*s/L*=0.2), whereas the amplitude of tail (*s/L*=0.8) and body (*s/L*=0.5) portions are less when compared to that of the curvature amplitude at the head portion in both kinds. A slight decrease in the amplitudes of the curvature at the body and tail portions observed in the later stages might be due to the increase in head amplitude (since the graph is normalised with the maximum head amplitude). Figure 5 shows the mean normalised frequency spectrum of *k* (*s,t*) of the mutant nematodes at different body positions in each stage. A single frequency peak of *f* ≈ 1.80 ± 0.21 Hz in stage 1, 2.25 ± 0.18 Hz in stage 2 and 1.95 ± 0.25 Hz in stage 3 is observed in the frequency spectrum. Only a negligible difference in the beating frequency is observed between the wild-type and mutant nematodes. Further, the variation observed in the bending curvature values 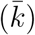 at the tail portion of the matured mutant nematodes did not significantly affect the beating frequency.

Figure 6 (*a*) shows the head beating amplitude (*A*_*h*_ in *mm*, corresponding to the peak in the Fourier spectrum) of both kinds of nematodes in each stage. The dashed lines (red and green) show that the *A*_*h*_ scales linearly with size for both kinds of nematodes investigated (mutant and wild-type) and the least square fit slope to the data yields the values of 0.49 (wild-type) and 0.41 (mutant). This slight difference in the measured slope values indicates that the *A*_*h*_ is modulated by the defective nature of the mutant nematodes as they mature. Also, based on the t-test, a significant difference in the mean head beating amplitude is observed between the wild-type and mutant nematodes in the later stages of their growth (as shown in figure 7 (*a*)).

**Figure 7:**
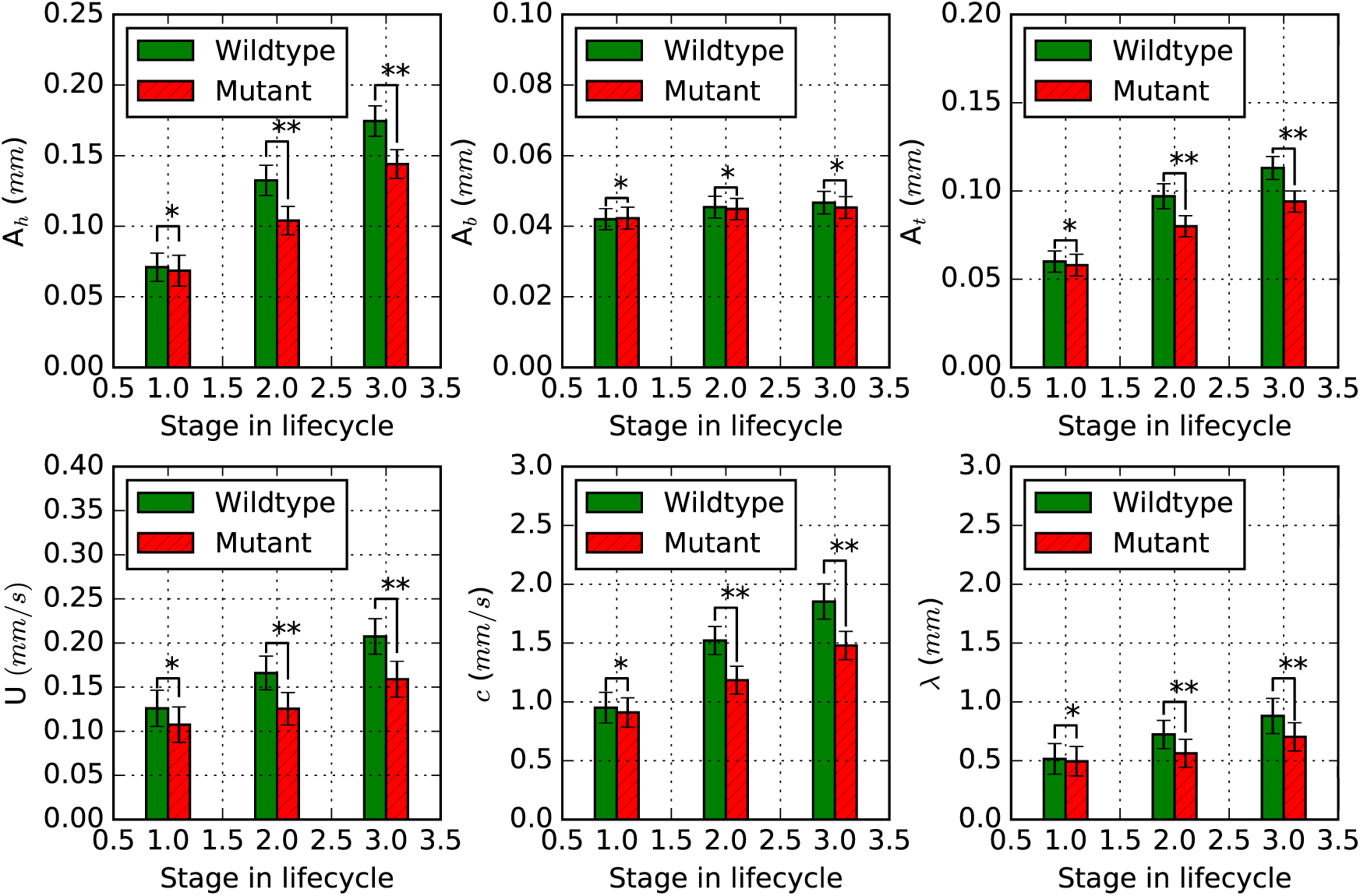
(*a*) Mean head beating amplitude (*A*_*h*_) at *s/L*=0.2. (*b*) Mean body beating amplitude (*A*_*b*_)at *s/L*=0.5. (*c*) Mean tail beating amplitude (*A*_*t*_) at *s/L*=0.8. (*d*) Mean swimming speed (*U*). (*e*) Mean wave speed (*c*). (*f*) Mean wave length (*λ*). Error bar represents the standard deviation of measured mean values in each stage of their growth. * denotes no significant difference and ** denotes significant difference based on t-test for p <0.05.

Figure 6 (*b*) shows the body beating amplitude (*A*_*b*_ in *mm*) of both kinds of nematodes in each stage of their growth. Only a slight increase in *A*_*b*_ is observed during their growth, and no significant difference in their magnitude is observed between wild-type and mutant nematodes. This observation suggests that *A*_*b*_ of mutant nematodes is not significantly modulated by the defective gene during their growth. Figure 6 (*c*) shows the tail beating amplitude (*A*_*t*_ in *mm*) of both kinds of nematodes in each stage. The *A*_*t*_ scales as a function of the size similar to *A*_*h*_ and the *A*_*t*_ of stage 2 and stage 3 mutants are found to be lower than that of the wildtype nematodes of the same size. The least square fit slope to the data yields the values of 0.24 (wild-type) and 0.19 (mutant), indicating that the *A*_*t*_ of the mutant nematodes are less than that of the wild-type nematodes as they grow. It is also worth noting that the larger values of *k* (*s,t*) estimated at the tail portion of the mutant nematodes in the later stages of their growth (as shown in figure 3) correlates with the decrease in *A*_*t*_ as they mature. This variation observed in *A*_*t*_ between the wild-type and mutant nematodes in their later stages as seen in figure 7 (*c*) (based on t-test) suggests that *A*_*t*_ is modulated by the effect of defective gene combined with ageing. This variation observed in the curvature values, head beating amplitude and tail beating amplitude of the matured mutant nematodes are found to be consistent with the observations of Sznitman et al. [2010b] where they investigated the kinematic properties of an adult *dys-1* mutant and showed much smaller beating amplitudes at the head and tail portion with increased curvature values (*k*).

Figure 6 (*d*) shows the effect of size and mutation on the swimming speed (*U*). Figure 7 (*d*) shows the average swimming speed of both kinds of nematodes computed in each stage. The mean swimming speed of the mutant nematodes in stage 1 shows a negligible difference when compared to the wild-type nematodes of the same size. However, significant variation is observed in the measured mean values between wild-type and mutant nematodes in their later stages, revealing that the swimming speed is modulated by the effect of the defective gene. It should also be noted that a decrease in the beating amplitude would also result in decreased swimming speed. Hence, the variation in mean *A*_*h*_ and *A*_*t*_ observed among the mutant nematodes as they mature is accompanied by the changes observed in the mean swimming speed.

Figure 6 (*f*) shows the effect of the defective gene on the wavelength (*λ*) during their growth. Similar to swimming speed and beating amplitude, the wavelength also scales linearly with size in both cases, and the least square fit slope to the data yields the values of 1.66 (wild-type) and 1.32 (mutant). Figure 7 (*f*) shows the mean wavelength of nematodes in each stage during their growth. The wavelength of the mutant nematodes in stage 1 shows a negligible difference when compared to that of the wild-type nematodes of the same size. However, similar to the mean swimming speed and mean beating amplitudes (*A*_*h*_ and *A*_*t*_), a significant difference is observed in the mean wavelength of the mutant nematodes as they mature.

### 3.3 Relationship between kinematic parameters

Figure 8 (*a*) shows the relationship between the swimming speed (*U*) and the velocity of neuromuscular wave (or) wave speed (*c*) in each stage. The swimming speed of both kinds of nematodes scales linearly with the velocity of the neuromuscular wave as they mature with conserved ratios of *U* /(*λf*) ≈ 1/7 (wild-type) and 1/9 (mutant). The ratios are much less than 1 in both cases and imply that the net swimming speed of both kinds of nematodes is much less than that of the velocity of the neuromuscular wave (wave speed, which is nearly twice that of the wavelength for a given frequency of ≈ 1.8-2.1 Hz. The variation observed in this ratio between wild-type and mutant nematodes is due to the decreased swimming speed and wavelength observed in their later stages.

**Figure 8:**
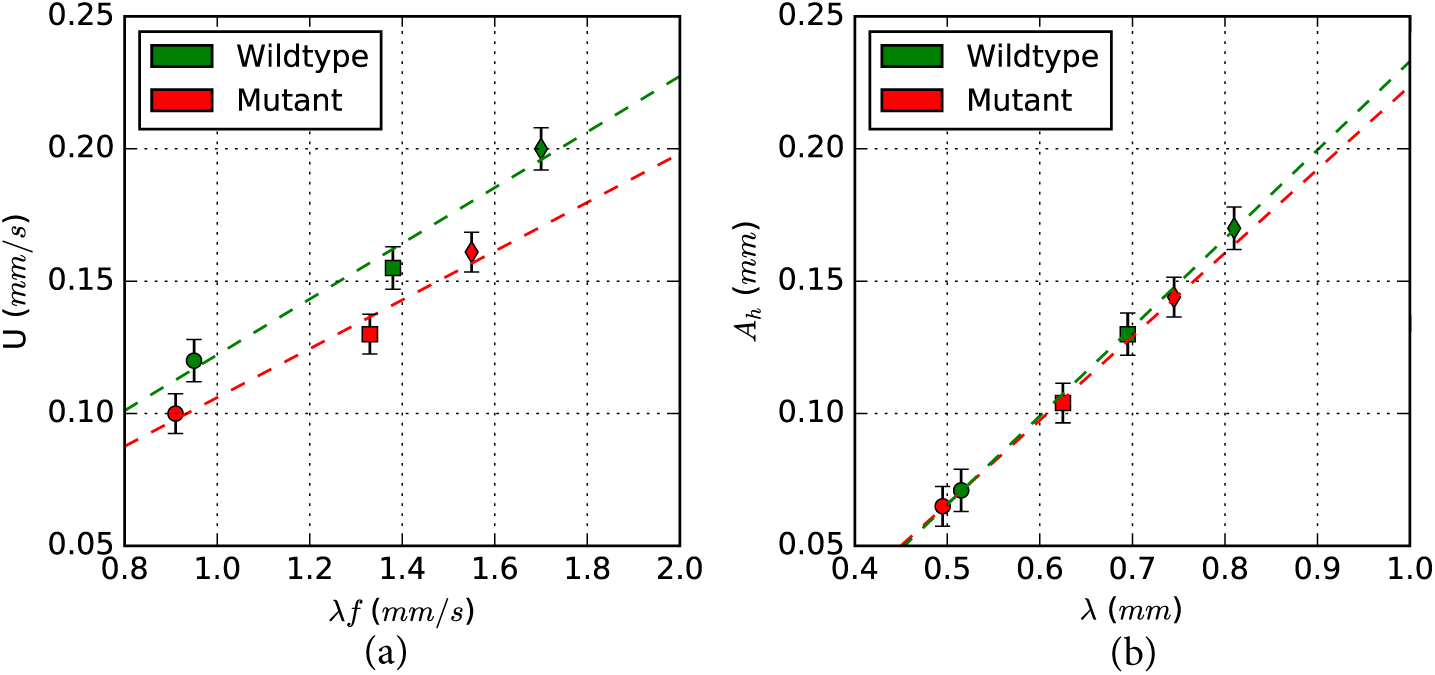
(*a*) Linear scaling of the swimming speed (*U*) with the wave speed (*c*) or the velocity of the neuromuscular waves. (*b*) Linear scaling of the head beating amplitude (*A*_*h*_) with the wavelength (*λ*). The symbols ∘, □ and ◊ represent the measured values in stage 1, stage 2 and stage 3 of the nematodes growth. The dashed lines represent the least square fit to the data.

Figure 8 (*b*) shows the linear scaling of the mean head beating amplitude (*A*_*h*_) with the wavelength (*λ*) in different stages of their growth. As the nematodes grow, the beating amplitude and wavelength of both kinds of nematodes increase in a correlated manner. The linear scaling of *A*_*h*_ with *λ* is almost similar in both cases and establishes a relationship, that is the least square fit to the kinematic data yields the regression lines *A*_*h*_ = 0.334*λ*-0.10 (wild-type) with a correlation coefficient (*R*^2^) of 0.98 and *A*_*h*_ = 0.315*λ*-0.091 (mutant) with a correlation coefficient (*R*^2^) of 0.99. The linear scaling also implies that the ratios (*A*_*h*_/*λ* ≈ 1/5 (wild-type) and 1/7 (mutant)) are conserved in both cases during their growth. This slight variation observed in the ratios between wild-type and mutant nematodes (as seen in the figure 8 (*b*)) is due to the decreased beating amplitude and wave length observed in their later stages, which further reveals that mutant exhibit motility defect as they mature.

### 3.4 Non-dimensional shape factor and swimming efficiency

The shape factor (*s*_*f*_) which is one of the properties of propagating waveform is defined as the ratio of wavelength (*λ*) to the length of the nematodes (*L*) [Gray and Lissmann, 1964]. Figure 9 (*a*) shows the *s*_*f*_ of both wild-type and mutant nematodes in each stage of their growth. It was observed that the *s*_*f*_ of swimming wild-type nematodes is a conserved ratio during their growth. This observation is consistent with the results of Karbowski et al. [2006], where they investigated the shape factor of the crawling wild-type nematodes and showed highly conserved ratios during their growth. Even though the ratios are conserved, the measured values of the *s*_*f*_ of the mutant nematodes are slightly lesser than that of the wild-type nematodes in the later stages of their growth due to the difference observed in the wavelength of the matured mutant nematodes (section 4.2.4), revealing the effect of defective gene on the swimming behaviour.

**Figure 9:**
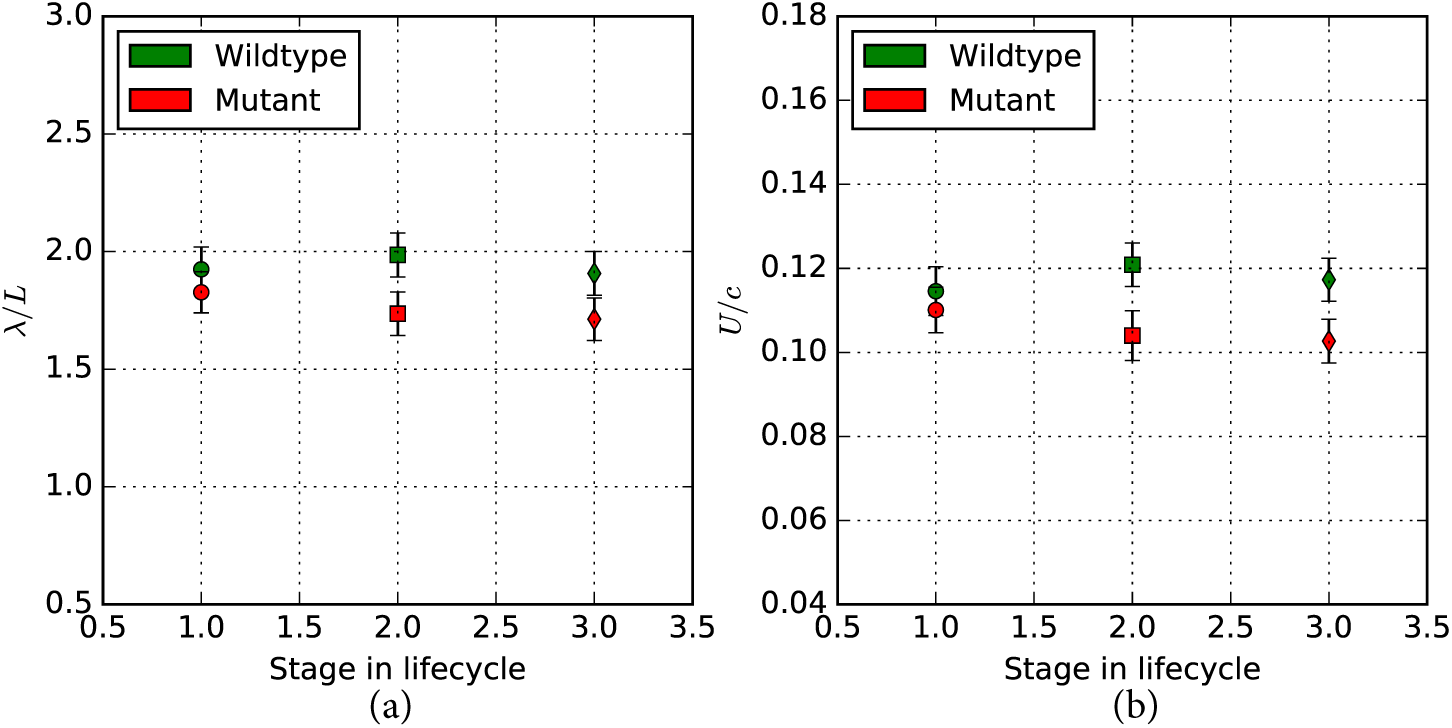
(*a*) Shape factor (*λ/L*) and (*b*) Swimming efficiency (*U/c*) in each stage of their growth. The symbols ∘, □ and ◊ represent the mean values in stage 1, stage 2 and stage 3 of the nematodes growth.

The swimming efficiency (*η*) is defined as a ratio of the swimming speed (*U*) to the wave speed (*c*). Figure 9 (*b*) shows the swimming efficiency of both wild-type and mutant nematodes in each stage of their growth. The swimming efficiency of the wild-type and mutant nematodes in the Newtonian fluid show negligible variation (slight increase) with size. The measured values of swimming efficiency of 0.114, 0.121 and 0.119 (wild-type), and 0.109, 0.106 and 0.102 (mutant) in the first three larval stages of their growth are in the same order of magnitude and consistent with the results (*η* ≈ 0.09-0.12) from previous experimental investigations of the adult wild-type nematodes (≈ 1 *mm* in length) [Gagnon et al., 2014, Sznitman et al., 2010d] swimming in fluids. These values measured are much less than one since their wave speed is higher than the swimming speed which in-turn is because of the fact that the nematodes slip during their undulating motion [Gray and Lissmann, 1964, Karbowski et al., 2006]. It was also observed that swimming efficiency of the mutant nematodes are lower than that of the wild-type nematodes in the later stages of their growth, revealing that the mutant investigated in this study is an inactive phenotype when compared to the swimming characteristics of the wild-type nematodes.

### 3.5 Fluid velocities and characteristics

Figure 10 shows the instantaneous velocity field generated by one of the sample wild-type nematode corresponding to the stage 1 in their life cycle. The colour code in the flow field illustrates the velocity magnitude of the fluid flow, and the arrows indicate the directions of fluid velocity at each point. Figure 10 shows the existence of small recirculation regions (vortices) along the body during swimming. These vortices are always attached to the body and flow in the opposite directions to each other. The existence of these vortices was confirmed throughout the swimming cycle, but the location of these vortices varies. These patterns observed are similar to the observations of Gray and Lissmann [1964], where they associated the maximum transverse displacement of nematode’s body to the local vortices (or) circulation regions attached to the nematode’s body. Previous investigations have also reported that three to four recirculation regions are observed along the adult nematodes body (≅1 mm in length) [Gray and Lissmann, 1964, Sznitman et al., 2010c] and pairs of adjacent vortices move in opposite directions to each other.

**Figure 10:**
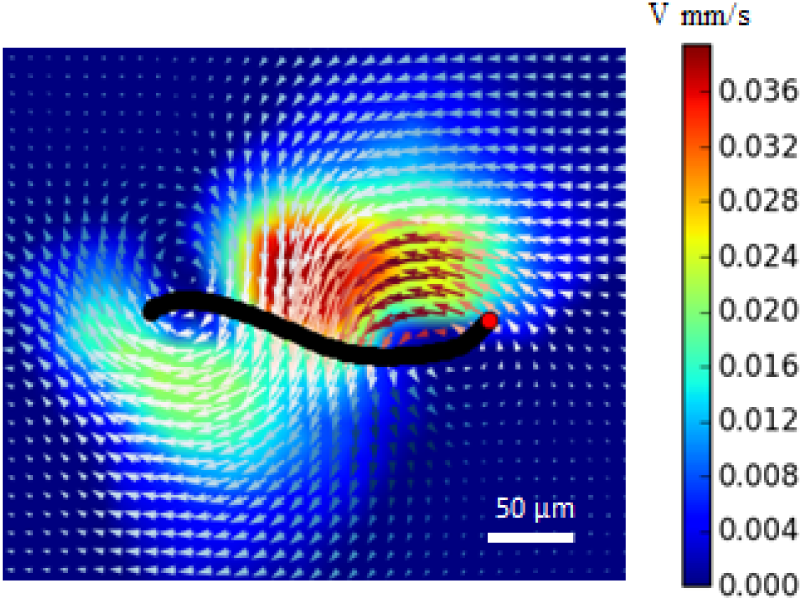
Velocity field generated by a sample wild-type nematode from stage 1.

Figure 11 and figure 12 show the instantaneous velocity fields of one of the sample wild-type and mutant nematodes in the fluid at different time intervals (t=0, T/5, 2T/5, and 3T/5, where T is the period of one beating cycle (T ≅ 0.5s)). The displacement of particles is maximum near the nematode’s body and decays rapidly away from the body, and far away from the nematode, the velocity magnitude is almost zero. It was found that the fluid velocity magnitude is very small (0.2-0.4 mm/s) and it undulates with respect to time. Also, the velocity magnitude and the features observed are consistent with the previous experimental studies of the adult wild-type nematode [Gray and Lissmann, 1964, Korta et al., 2007, Sznitman et al., 2010c].

**Figure 11:**
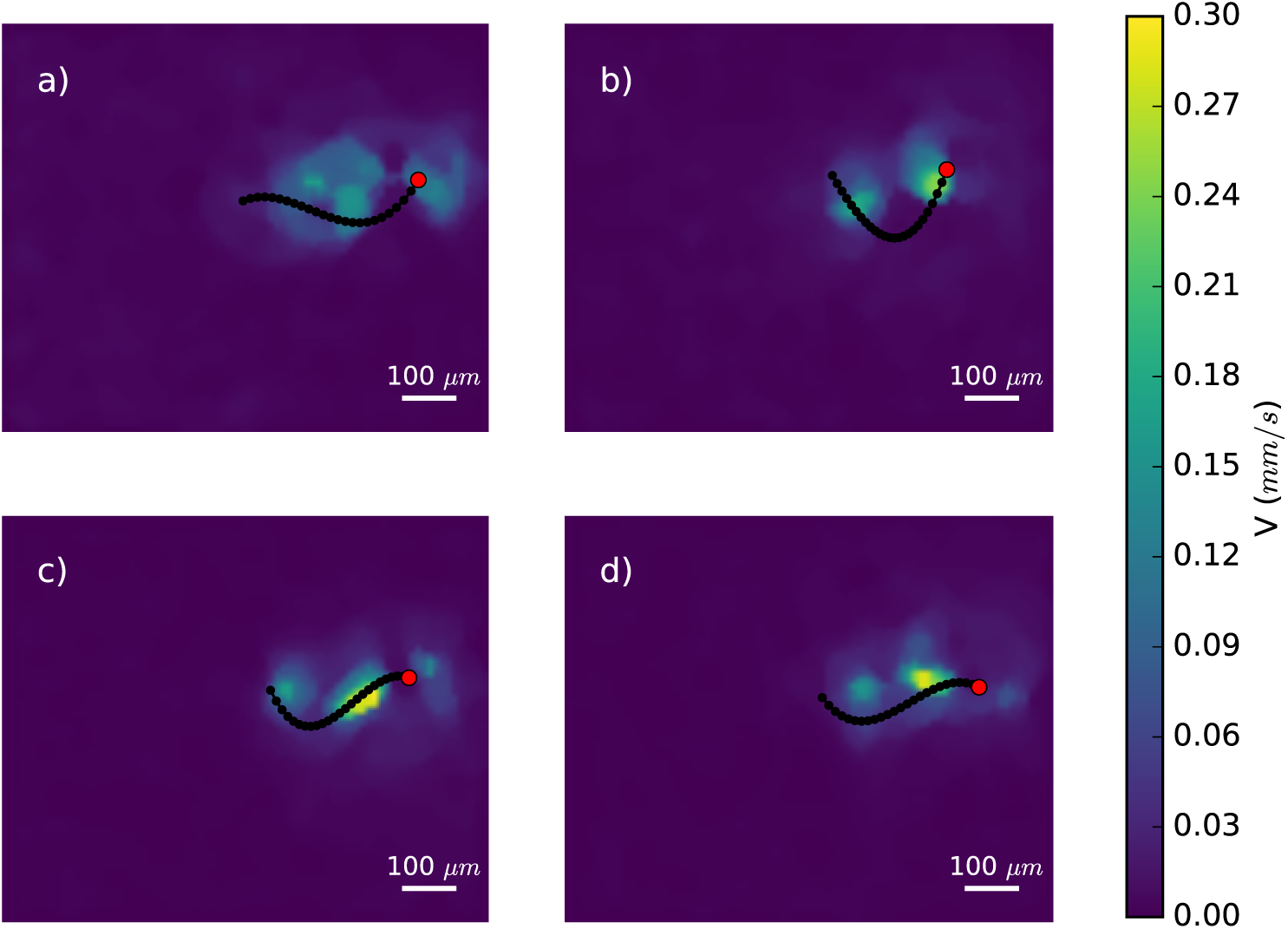
(*a, b, c, d*) Fluid velocity magnitude corresponding to a sample wild-type nematode at different time steps (t=0, T/5, 2T/5, and 3T/5), where T is the period of one beating cycle (≈ 0.5s).

**Figure 12:**
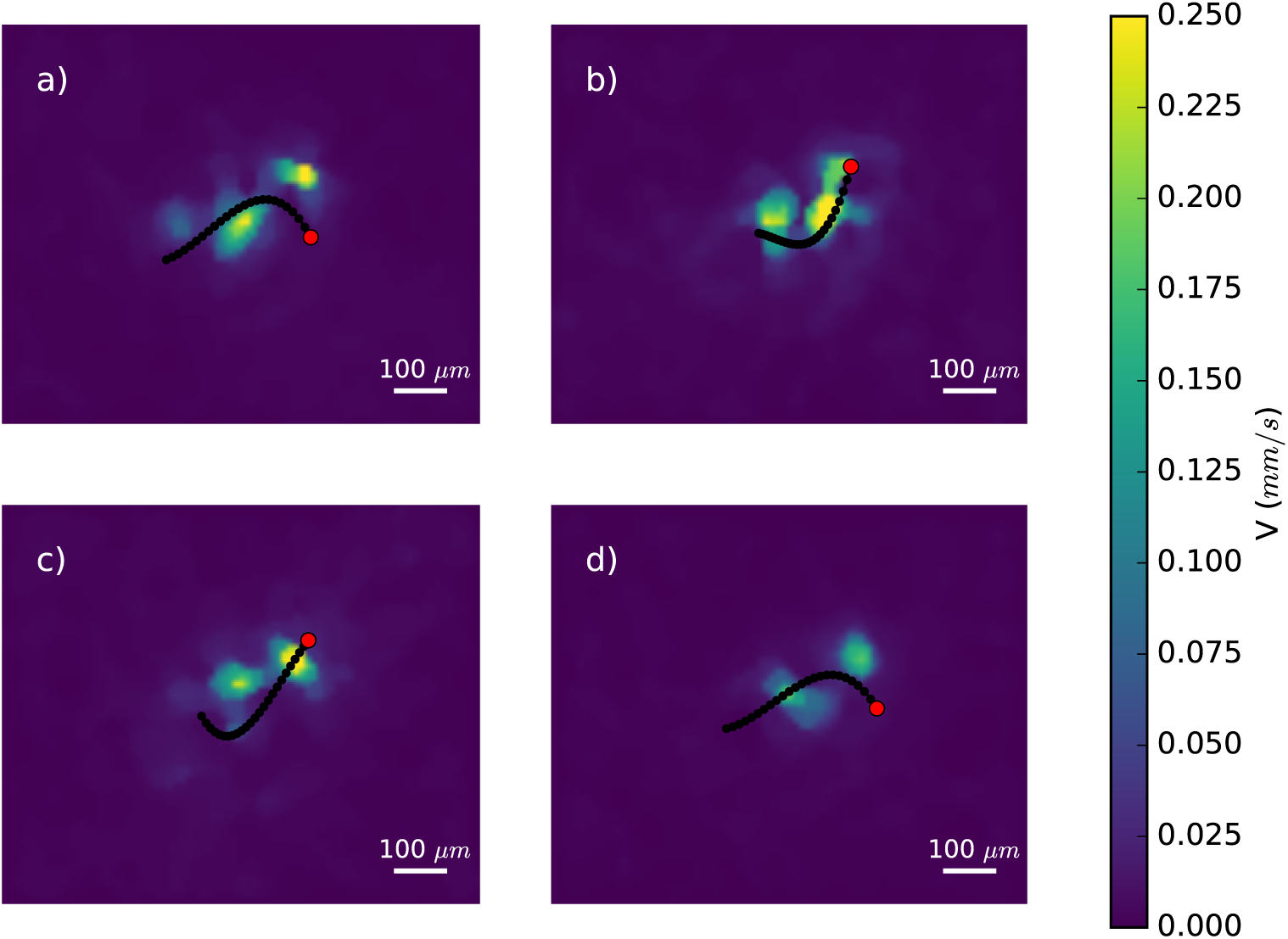
(*a, b, c, d*) Fluid velocity magnitude corresponding to a sample mutant nematode at different time steps (t=0, T/5, 2T/5, and 3T/5), where T is the period of one beating cycle (≈ 0.5s).

### 3.6 Power exerted by the wild-type and mutant nematodes

Figure 13 shows the estimated power (*P*) based on the limited 2C-2D PIV measurements in the mid-plane of the nematode exerted by individual nematode samples computed in each stage of their growth. The data show that the power exerted by the nematodes in Newtonian fluid increases linearly with size, indicating that the motility is modulated by the nematodes growth. The estimated magnitude of power measured in this study and the estimates from previous experimental investigations of the adult wild-type nematodes in Newtonian fluids are in the same order of magnitude [Sznitman et al., 2010c, Krajacic et al., 2012, Wan et al., 2014]. Figure 13 (*b*) shows the estimated mean power 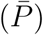 exerted by the nematodes in each stage of their growth. The estimated power exerted by the wild-type and mutant nematodes showed negligible difference in stage 1. A significant difference is observed in the estimated power exerted by the mutant nematodes as they mature (S2 ≈ 1.06 ± 0.333, S3 ≈ 2.1 ± 0.37 *pW/mm*) when compared to that of the wild-type nematodes (S2 ≈ 2.2 ± 0.35, S3 ≈ 3.65 ± 0.359 *pW/mm*) of the same size. This variation in estimated power exerted by the mutant nematodes in their later stages clearly shows that the defective gene modulates the power exerted by the mutants. This variation is also accompanied by the changes observed in the primary kinematic parameters including the beating amplitude, wavelength and swimming speed of the mutant nematodes in their later stages, revealing that the matured mutant nematodes are less active than the wild-type nematodes of the same size. Based on these results, the mutant with motility defect investigated in this study is classified as a less active (inactive) phenotype.

**Figure 13:**
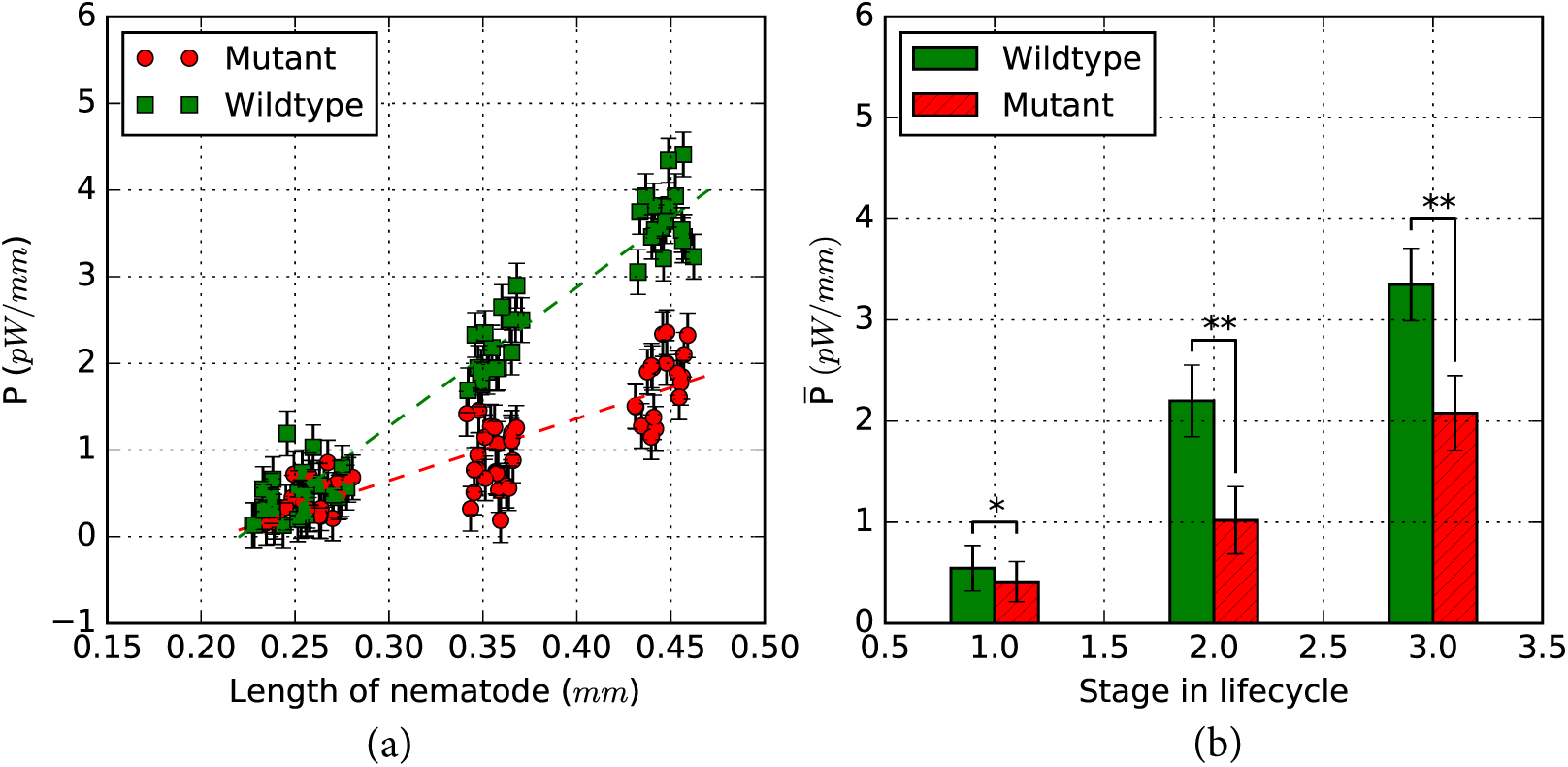
(*a*) Power (*P*) exerted by wild-type and mutant nematodes in the stationary fluid medium. Error bar represents the standard error across samples. The dashed lines represent the linear fit to the data. (*b*) Mean power exerted 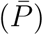 by wild-type and mutant nematodes in the stationary fluid medium in each stage. Error bar represents the standard deviation. * denotes no significant difference and ** denotes significant difference based on t-test for p <0.05.

## 4 Conclusions

The present study investigated the swimming characteristics of the wild-type and *m*ir-1 mutant nematodes at each stage of their growth using 2C-2D *µ*-PIV. The kinematic study of both nematodes strains during their growth provided novel insight by providing quantitative kinematic data such as the bending curvature (*k*), frequency of beating (*f*), amplitude of beating (*A*) and wavelength (*λ*) as a function of size of the nematodes.

The beating frequency of the wild-type and mutant nematodes is not significantly influenced by their growth. The temporal form of the swimming gait is also found to be highly robust, independent of body positions and not affected by the size. The swimming speed, beating amplitude (head and tail), wavelength and power exerted by both kinds of nematodes are found to scale linearly with size with the ratios *A*_*h*_/*λ* and *U* /(*λf*) found to be are conserved during their growth.

Mutant nematodes (in larval stages 2 and 3) move slower than the wild-type nematodes of the same size with significant difference in the beating amplitude (head and tail) and wavelength of the mutant nematodes, revealing that the mutant exhibits motility defect. The swimming efficiency and the shape factor of the matured mutant nematodes are slightly lower than that of the wild-type nematodes of the same size, revealing that the mutant with motility defect shows behavioural variations as a result of ageing.

From the fluid flow measurements it is found that the *m*ir-1 mutant investigated in this study exerts less power in their later stages of aging when compared to that of the wild-type nematodes, again revealing the age-related effect of the mutation in regulating the swimming behaviour. Furthermore, there is a significant variation in the magnitude of local circulation of fluid around the mutant nematodes, indicating that the flow field is modulated by the effect of the mutation. Based on these results (*i*.e. the observations from the kinematic study and power exerted), the mutant investigated in this study is classified as a less active (*i*.e. in-active) phenotype when compared to that of the wild-type nematode.

This study has demonstrated that high-spatial resolution simultaneous imaging of the motion of *C.elegans* in a fluid seeded with tracer particles using a 2C-2D *µ*-PIV imaging set-up can provide detailed kinematic measurements of the nematodes with simultaneous fluid flow measurements as a function of their age. The latter in addition to providing quantitative data of the fluid flow and hence, the swimming gait of the nematodes, also provides detailed quantitative energy measurements during the life of the nematodes, which is a direct measurement of activity of the nematodes via its energy exertion as function of its age.

## 5 Acknowledgement

We would like to acknowledge Leelee Ng from Biomedicine Discovery Institute at Monash University for their technical support and advice on *C. elegans*. We would also like to acknowledge the funding provided by a Monash IDR grant to undertake this research and equipment and facilities funded through ARC LIEF grants that supported this project.

## References

M. Backholm, A. K. S. Kasper, R. D. Schulman, W. S. Ryu, and K. D. Veress. The effects of viscosity on the undulatory swimming dynamics of *C. elegans*. Physics of Fluids, 27:091901, 2015.

S. Brenner. The genetics of *C.elegans*. Genetics, 77:71–94, 1974.

N. Chen, T. W. Harris, I. Antoshechkin, C. Bastiani, and et al. WormBase: a comprehensive data resource for Caenorhabditis biology and genomics. Nucleic Acids Research, 33:D383–D389, 2005.

C. J. Cronin, J. E. Mendel, S. Mukthar, Y. M. Kim, and R. C. Stirb. An automated system for measuring the parameters of nematode sinusoidal movement. BMC Genetics, 6:5, 2005.

M. Fedrizzi and J. Soria. Application of a single-board computer as a low cost pulse generator. Measurement Science and Technology, 26:095302, 2015.

D. A. Gagnon and P. Arratia. The cost of swimming in generalized Newtonian fluids: experiments with *C.elegans*. Journal of Fluid Mechanics, 800:753–765, 2016.

D. A. Gagnon, N. C. Keim, and P. E. Arratia. Undulatory swimming in shear-thinning fluids: experiments with *Caenorhabditis elegans*. Journal of Fluid Mechanics, 758:1–11, 2014.

J. Gray and G. J. Hancock. The propulsion of sea urchin spermatozoa. Journal of Experimental Biology, 32:2814, 1955.

J. Gray and H. W. Lissmann. The locomotion of nematodes. Journal of Experimental Biology, 41:135, 1964.

I. K. Hariharan, Haber, and A. Daniel. Yeast, Flies, Worms, and Fish in the Study of Human Disease. The New England Journal of Medicine, 348:2457–2463, 2003.

T. W. Harris, N. Chen, F. Cunningham, and M. Tello-Ruiz. Worm Base, a multispecies resource for nematode biology and genomics. Nucleic Acids Research, 32:411–417, 2004.

D. Hart. The elimination of correlation error in PIV processing. In 9^th^ International Symposium of Applications of Laser Techniques to Fluid Mechanics, pages I:13.3.1 – 13.3.8, Lisbon, Portugal, July 1998.

Z. Hu, S. Hom, T. Kudze, X.-J. Tong, S. Choi, G. Aramuni, W. Zhang, and J. M. Kaplan. Neurexin and neuroligin mediate retrograde synaptic inhibition in *C. elegans*. Science, 337: 980–984, 2012.

T. D. M. Johnson, D. A. Gagnon, P. E. Arratia, and E. Lauga. Flow analysis of the low Reynolds number swimmer *C.elegans*. Physical Review Fluids, 1:1–9, 2016.

J. Karbowski, C. J. Cronin, A. Seah, J. E. Mendel, D. Cleary, and P. W. Sternberg. Conservation rules, their breakdown, and optimality in *Caenorhabditis elegans* sinusoidal locomotion. Journal of Theoretical Biology, 242:652 – 669, 2006.

J. Korta, D. A. Clark, C. V. Gabel, L. Mahadevan, and A. D. T. Samuel. Mechanosensation and mechanical load modulate the locomotory gait of swimming *C.elegans*. Journal of Experimental Biology, 210:2383–9, 2007.

P. Krajacic, X. N. Shen, P. K. Purohit, P. E. Arratia, and T. Lamitina. Biomechanical profiling of *Caenorhabditis elegans* motility. Genetics, 191:1015–1021, 2012.

L. Lam, S. W. Lee, and C. Y. Suen. Thinning Methodologies-A Comprehensive Survey. IEEE Transactions on Pattern Analysis and Machine Intelligence, 14:869–885, 1992.

W. Li, Z. Feng, P. Sternberg, and X. Xu. A *C.elegans* stretch receptor neuron revealed by a mechanosensitive TRP channel homologue. Nature, 440:684–687, 2006.

J. Lighthill. Flagellar hydrodynamics. SIAM Rev., 18:161–230, 1976.

M. Maria and T. Nektarios. Modelling human diseases in *Caenorhabditis elegans*. Biotechnology Journal, 5:1261–1276, 2010.

S. Ongwattanakul, P. Chewputtanagul, D. J. Jackson, and K. G. Ricks. Scalable Giga-Pixel binary image morphological operations. Proceedings of the 2003 International Symposium on Circuits and System, 2:09170, 2003.

A. M. Persico and V. Napolioni. Autism genetics. Behav. Brain Res., 251:95 – 112, 2013.

E. M. Purcell. Life at low Reynolds Number. American Journal of Physics, 45:3–11, 1977.

S. Ravikumar. Experimental investigation of the motility of wildtype and mutant caenorhabditis elegans using *µ*-piv. Master’s thesis, Monash University, Melbourne, Victoria, Australia, 7 2018.

X. Shen, J. Sznitman, P. Krajacic, T. Lamitina, and P. E. Arratia. Undulatory locomotion of *C. elegans* on wet surfaces. Biophysical Journal, 102:2772–2781, 2012.

D. J. Simon, J. M. Madison, A. L. Conery, K. L. Thompson-Peer, M. Soskis, G. B. Ruvkun, J. M. Kaplan, and J. K. Kim. The microrna mir-1 regulates a mef-2-dependent retrograde signal at neuromuscular junctions. Cell, 33(5):903 – 915, 2008. ISSN 0092-8674. doi: https://doi.org/10.1016/j.cell.2008.04.035. URL http://www.sciencedirect.com/science/article/pii/S0092867408006090.

P. Soille. Morphological Image Analysis: Principles and Applications. Springer, 14, 1999.

J. Soria. Digital cross-correlation particle image velocimetry measurements in the near wake of a circular cylinder. In Int. Colloquium on Jets, Wakes and Shear Layers, pages 25.1 – 25.8, Melbourne, Australia, 1994. CSIRO.

J. Soria. An investigation of near wake of a circular cylinder using a video-based digital cross correlation particle image velocimetry technique. Experimental Thermal and Fluid Science, 12:221–233, 1996.

J. Soria. Multigrid approach to cross-correlation digital PIV and HPIV analysis. In 13th Australasian Fluid Mechanics Conference, pages 381–384, Monash University, Melbourne, 1998.

H. A. Stone and A. D. T. Samuel. Propulsion of Microorganisms by Surface Distortions. Physical Review Letters, 77:4102–4104, 1996.

J. Sznitman, P. K. Purohit, P. Krajacic, T. Lamitina, and P. E. Arratia. The effects of fluid viscosity on the kinematics and material properties of *C.elegans* swimming at low Reynolds number. Experimental Mechanics, 50:1303–1311, 2010a.

J. Sznitman, P. K. Purohit, P. Krajacic, T. Lamitina, and P. E. Arratia. Material Properties of *Caenorhabditis elegans* Swimming at Low Reynolds Number. Biophysical Journal, 98: 617–626, 2010b.

J. Sznitman, X. Shen, R. Sznitman, and P. E. Arratia. Propulsive force measurements and flow behaviour of undulatory swimmers at low Reynolds number. Reports on Progress in Physics, 22:121901, 2010c.

R. Sznitman, M. Gupta, G. D. Hager, P. E. Arratia, and J. Sznitman. Multi-environment model estimation for motility analysis of *Caenorhabditis elegans*. Plos One, 5:e11631, 2010d.

N. Tavernarakis, W. Shreffler, S. L. Wang, and M. Driscoll. unc8, a DEG/ENaC family member, encodes a subunit of a candidate mechanically gated channel that modulates C. elegans locomotion. Neuron, 18:107–119, 1997.

R. van den Boomgard and R. van Balen. Methods for Fast Morphological Image Transforms Using Bitmapped Images. Computer Vision, Graphics, and Image Processing: Graphical Models and Image Processing, 54:254–258, 1992.

K. von Ellenrieder, J. Kostas, and J. Soria. Measurements of a wall-bounded, turbulent, separated flow using hpiv. Journal of Turbulence - selected papers from the 8th European Turbulence Conference, 2001.

J. K. Wan, S. S. Yue, and S. C. Han. Characteristics of kinetic power and propulsion of the nematode *C.elegans* based on a micro-particle image velocimetry system. Biomicrofluidics, 8:024116, 2014.

J. Westerweel and F. Scarano. Universal outlier detection for PIV data. Experiments in Fluids, 39:1096–1100, 2005.

